# QuAdTrim: Overcoming computational bottlenecks in sequence quality control

**DOI:** 10.1101/2019.12.18.870642

**Authors:** Andrew J. Robinson, Elizabeth M. Ross

## Abstract

With the recent torrent of high throughput sequencing (HTS) data the necessity for highly efficient algorithms for common tasks is paramount. One task for which the basis for all further analysis of HTS data is initial data quality control, that is, the removal or trimming of poor quality reads from the dataset. Here we present QuAdTrim, a quality control and adapter trimming algorithm for HTS data that is up to 57 times faster and uses less than 0.06% of the memory of other commonly used HTS quality control programs. QuAdTrim will reduce the time and memory required for quality control of HTS data, and in doing, will reduce the computational demands of a fundamental step in HTS data analysis. Additionally, QuAdTrim impliments the removal of homopolymer Gs from the 3’ end of sequence reads, a common error generated on the NovaSeq, NextSeq and iSeq100 platforms.

**Availability and Implementation:** The source code is freely available on bitbucket under a BSD licence, see COPYING file for details: https://bitbucket.org/arobinson/quadtrim

**Contact:** Andrew Robinson andrewjrobinson at gmail dot com

## Introduction

In recent years the amount of high throughput sequencing data being generated has skyrocketed, and this trend is set to continue. In bioinformatics, the saying ‘Garbage in, garbage out’ still rings true implying that the quality of data used in analyses is of utmost importance to the integrity of the results. Fundamentally, the first step in data analysis in any HTS project is quality control of the sequence reads. A number of common tools are available for this task (Bolger, et al., 2014; Joshi and Fass, 2011; Martin, 2011; Shrestha, et al., 2014), however these tools fall short of current requirement either due to long processing times, larger memory usage, or unexpected behaviour. As such it is timely for the development of new algorithms that focus both on efficiency (due to the large amount of data being generated, and the likelihood that the data deluge will continue to increase for several years to come) and accuracy of results.

## Methods

### Algorithm

The current version (v2.0.1) of the QuAdTrim algorithm is described. The algorithm was implemented in C++ and has two main phases (Figure 1):

- Adapter: removes overhanging adapters (for when there are short insert sizes)
  - it has the option to collect the adapters that were trimmed so that they can be used to search for failed reads that consisted of entirely adapter sequence
- Quality: removes bases from ends of sequence in two modes:
  - 3’ only: typical usage for Illumina reads
    - Additionally erroneous tailing G’s can be removed, typical of NextSeq, iSeq100 and NovaSeq reads with short or no insert.
  - 5’ and 3’: remove bases from both ends of reads
    - Recovering data from runs with interruptions etc.

**Figure 1.**
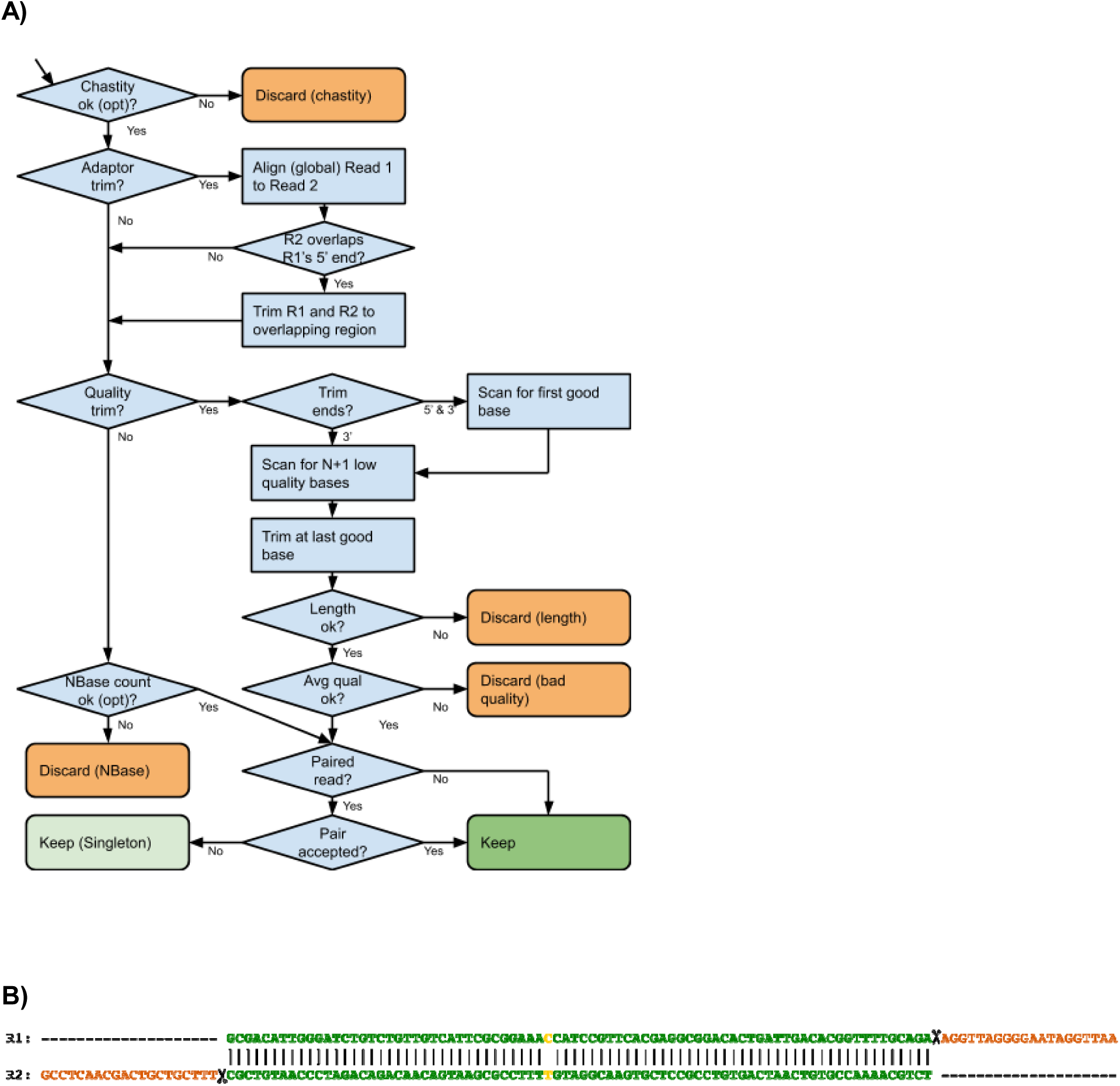
QuAdTrim algorithm description. **A)** Flowchart depicting the algorithm of the QuAdTrim application for each sequence (or pair of sequences). **B)** An example alignment of two reads highlighting the adapter trimming process. Read 2 (R2) is shown in reverse order to what it appears in the file so that its bases line up with read 1. The scissor icon highlights the point where the reads will be trimmed. The orange bases at the end of each read are the adaptor sequence that will be removed from the output. The dashes that match are placeholders to line up the sequences visually and are not present in the input or output files.

Additionally QuAdTrim has two ancillary phases. These are performed alongside the main two phases

- Average Quality:
  - If the average quality of the final trimmed read is below a provided threshold then it is discarded.
- Ambiguous Base count:
  - If there are more than a given number of ambiguous bases in the read then it will be discarded.

### Tool Comparison and raw data

The performance of QuAdTrim was compared to Trimmomatic, CutAdapt, QTrim and Sickle. Where possible, parameters were set to the same values (thresholds) to perform comparisons between tools.

For base pair quality control on single ended Illumina data the tool parameters were: QuAdTrim: quadtrim -m 2 -q 5 -a 1 test1.fq

Trimmomatic: java -jar trimmomatic-0.32.jar SE -threads 1 -phred33 test1.fq test1-pass.fq

TRAILING:20 SLIDINGWINDOW:50:20

QTrim: QTrim_v1_1.py -o output -fastq test1.fq

Sickle: sickle se -f test1.fq -t sanger -o test1-pass.fq -q 20 -l 50

For adapter contamination removal on paired end Illumina data Adapter Trimming the tool parameters were:

QuAdTrim: quadtrim -m 1 -T 0 -l 50 -S ACAGGCTGCGGAAATTACGTTAGTCCCGTCAGTAAAATTA -S TGGCACCTACACGGACCGCCGCCACCGCCGCGCCACCATA test1.fq test2.fq

Trimmomatic: java -jar trimmomatic-0.32.jar PE -threads 1 -phred33 test1.fq test2.fq test1-pass.fq test1-singleton.fq test2-pass.fq test2-singleton.fq ILLUMINACLIP:adapter.fa:2:30:10 MINLEN:50

Cutadapt: cutadapt -a ACAGGCTGCGGAAATTACGTTAGTCCCGTCAGTAAAATTA -a TGGCACCTACACGGACCGCCGCCACCGCCGCGCCACCATA --minimum-length 50 -- paired-output tmp.2.fastq -o tmp.1.fastq test1.fq test2.fq cutadapt -a ACAGGCTGCGGAAATTACGTTAGTCCCGTCAGTAAAATTA -a TGGCACCTACACGGACCGCCGCCACCGCCGCGCCACCATA --minimum-length 50 -- paired-output test1-pass.fq -o test2-pass.fq tmp.2.fastq tmp.1.fastq rm tmp.1.fastq tmp.2.fastq

### Memory and runtime comparison

Each tool was run 4 times back-to-back with a 1 second delay. The first run was discarded so that each of the 3 remaining runs could have a similar amount of system cache hits. The 4 runs for each tool were run back-to-back on the same machine with no other task executing to give an equal execution environment.

### Removal of Homopolymer suffix in NovaSeq, NextSeq and iSeq100 data

To illustrate the remove of homopolymer G’s that are induced by the one and two colour chemistry used on the NovaSeq, NextSeq and iSeq100 platforms, base calls from reads generated on the iSeq100 platform were trimmed with QuAdTrim with and without the ‘-g’ flag (q*uadtrim -m 2 -q 20 - a 20 -l 50 -p 3 iseq*.*fastq* OR *quadtrim -m 2 -q 20 -a 20 -l 50 -p 3 -g iseq*.*fastq*). As a control, the number of poly-C present at the end of the same set of sequence reads was also assessed. Because of the complimentary pairing of G-C, the number of true (present in the sequence genome, not those generated by sequencing artefacts) G homopolymers is expected to be approximated by the number of C homopolymers.

## Results and Discussion

To address the requirements of HTS we developed QuAdTrim, a quality control software designed for (but not limited to) Illumina data, with the ability to take both paired and single reads, with adjustable quality settings to enable the user to customise the quality control to suit their task. QuAdTrim is able to detect partial adapter sequences in paired read data by using reverse complement overlaps between pairs. These partial adapter sequences are caused by the DNA insert in the sequenced molecule being shorter than the read length, in which case the molecule is sequenced through the DNA and into the alternative end adapter and barcode sequence. Additionally, the algorithm is able to detect and remove erroneous repetitions of ‘G’ introduced at the end of NextSeq sequence reads.

### Benchmarking

We compared the performance of QuAdTrim with the commonly used Trimmomatic, Sickle and QTrim Bolger, et al. (2014); Martin (2011); Shrestha, et al. (2014). QuAdTrim had both the fastest runtimes and lowest memory usage of all programs tested (Figure 2). When trimming reads based on quality scores QuAdTrim showed a 34% and 73% improvement in running time compared to Trimmomatic and Sickle respectively, with 5651% improvement (i.e. 57 times faster) compared to QTrim. When used in adapter removal mode, QuAdTrim run time showed a 145% improvement compared to Trimmomatic’s adapter trimming performance, and an 897% improvement compared to cutadapt. The amount of memory used by Trimmomatic was substantially higher than all other programs in both quality trimming and adapter removal modes (Figure 2B). Both QuAdTrim and Sickle used substantially less memory than QTrim in quality trimming mode, and QuAdTrim showed a 1190% improvement compared to cutadapt in memory use.

**Figure 2.**
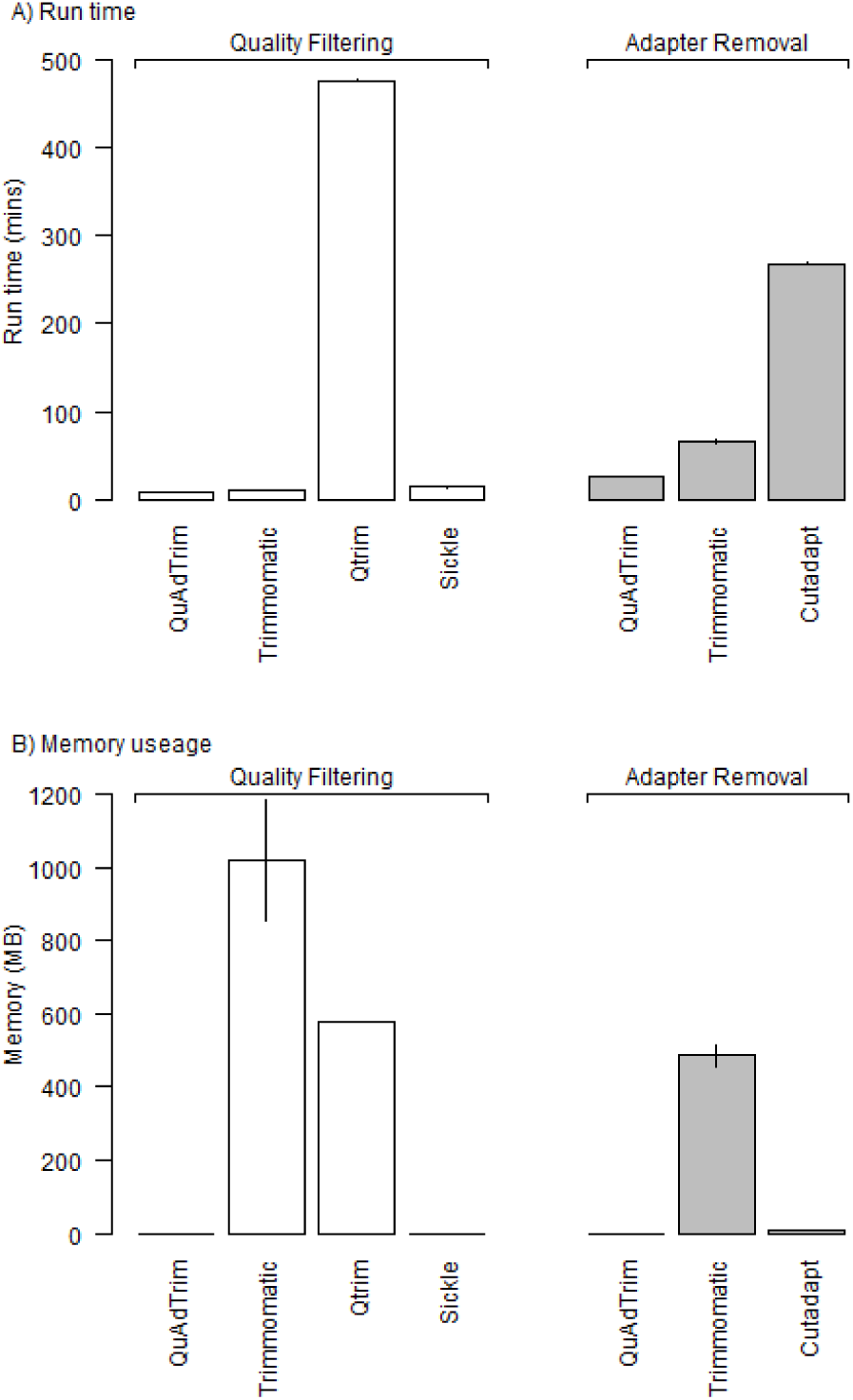
Computational time and memory usage for QuAdTrim and other popular software. Quality filtering: filtering of data based on Phred Q scores to remove low quality bases, data consisted of 1.7GB (4,759,058 reads of 151bp in length) single end Illumina HiSeq1500 data. Adapter filtering: removal of adapter or adapter-like sequences, data consisted of 3.4GB (4,759,058 pairs of 151bp in length) paired end Illumina HiSeq1500 data. Values and the average of three replicate runs (all single core with default/recommended parameters for each program), with error bars showing S.E.M.

To ensure the accuracy of the filtered data we compared the filtered output (i.e. reads that passed quality control) (Table 1). Trimmomatic and QuAdTrim had a high level of agreement, with only 4 reads being treated differently between the two algorithms. These reads were discarded by QuAdTrim but not Trimmomatic). QTrim also reported similar results to QuAdTrim and Trimmomatic, while Sickle showed substantially different treatment of read in comparison to QuAdTrim and the other popular programs tested.

**Table 1.** Pairwise comparison of quality filtering differences. Filtering was performed on data consisting of 1.7GB (4,759,058 reads of 151bp in length) single end Illumina HiSeq1500 data. The number of reads discarded by only one program is shown as a measure of disagreement between algorithms.

|  | QuAdTrim | Trimmomatic | QTrim |
| --- | --- | --- | --- |
| Trimmomatic | 4 | - | - |
| QTrim | 491 | 487 | - |
| Sickle | 35452 | 35448 | 35933 |

Adapter removal with QuAdTrim is sensitive for short adapter fragments, such as those less that 10bp, however long fragments, particularly those where no insert is present (i.e. two adapters are ligated directly to each other) cannot be identified with QuAdTrim due to the algorithms use of reverse compliment overlaps in identifying the beginning and ends of the DNA insert fragment. As such, for sequence libraries where adapters are present due to the lack of DNA insert, using a homology based identification algorithm such as cutadapt is recommended. However if adapter sequences are present due to short inserts, QuAdTrim has the advantage of being much faster than cutadapt, and also being implemented as part of the larger QC pipeline, thus removing the need to process data twice (once to remove low quality bases and once for adapter filtering). Additionally, QuAdTrim is able to report the sequences that have been identified as adapter, which can subsequently inform users on the adapter sequences used if this information is otherwise unavailable. Such knowledge is necessary for cutadapt and as such a two tier adapter removal methodology may be implemented: quality and adapter filtering with QuAdTrim, and then subsequent adapter identification with cutadapt using the sequences identified in the previous step.

Here we have presented QuAdTrim and compared it to quality control algorithms commonly used on HTS data. QuAdTrim maintains the accuracy of Trimmomatic, while significantly reducing the wall time and memory required. While QuAdTrim and sickle have similar computational resource demands, the output of sickle significantly differs from other trimming programs while QuAdTrim produces almost identical output.

### Homopolymer G removal

QuAdTrim has poly-G removal enabled. Taking the compliment of the ‘G’ base as a control (‘C’), the number of polyG of different lengths was compared between raw, trimmed with –g on and trimmed without –g were compared. Homoploymer Gs were abundant at the 3’ ends of the untrimmed data (Figure 3). Regardless of the presence of the –g flag, 123,320 reads (88.4%) passed the trimming criteria. Trimming without the –g flag reduced the number of reads with homopolymer G slightly. Trimming with the –g flag reduced the number of reads ending in homopolymer G dramatically, to below the number of homopolymer Cs. It is expected that the homopolymer C bases are an approximation of the true homopolymers in the sample, and thus the –g flag likely removes some true homoploymer Gs from the data. Thus, if users are interested in regions containing homopolymers it is advisable to consider the value of the –g flag. For most cases however, the –g flag successfully removes homopolymer G runs from the 5’ end of Illumina reads from the iSeq100 platform.

**Figure 3.**
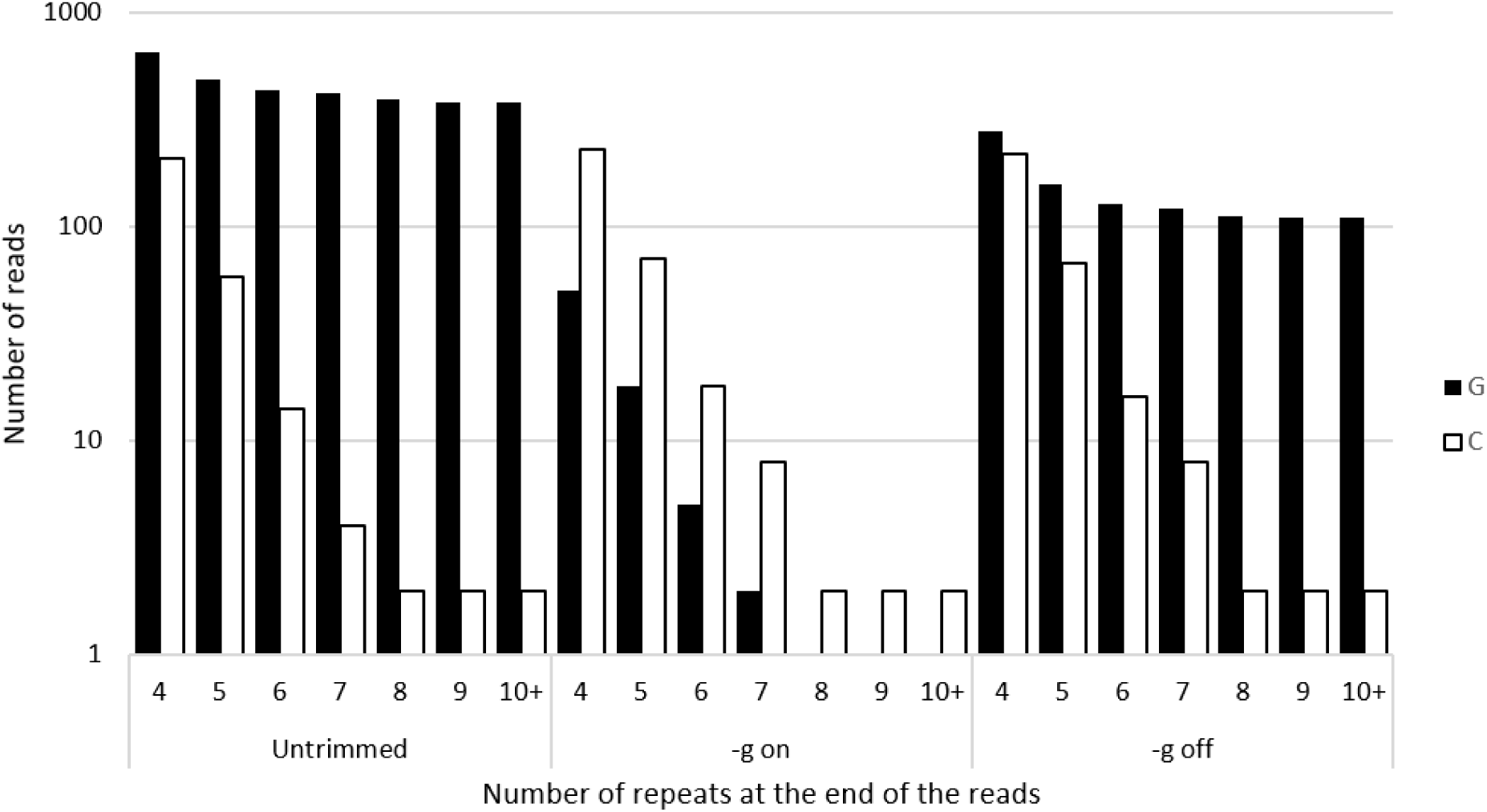
Number of reads from single end trimming of a DNA sample sequenced on the iSeq100. Total reads trimmed: 34884. The X axis indicates the number of homopolymer bases observed at the end of the reads pre- (untrimmed) and post-trimming, with (-g on) and without (-g off) the poly G removal enabled. For comparison the number of Poly-C at the end of the reads is included.

### History, usage and future directions

QuAdTrim was first developed in 2011 for use in a metagenomic study that utilised Illumina paired end sequence data (Ross, et al., 2012). Subsequently QuAdTrim has been used for quality control of Illumina sequence data in from human, bacterial, virus, cattle, sheep, and salmon studies amoung others (Bolormaa, et al., 2019; Daetwyler, et al., 2017; Kamato, et al., 2017; Kijas, et al., 2019; Kijas, et al., 2018; Ross, et al., 2013; Ross, et al., 2013; Wei, et al., 2018).

Currently the genomics community is experiencing an explosion of long read sequence data. QuAdTrim is currently under development to incorporate quality control of long read data so that that a single tool will be available for the quality control of both short and long read data.

## Acknowledgement

This work was supported by the VLSCI’s Life Sciences Computation Centre, a collaboration between Melbourne, Monash and La Trobe Universities and an initiative of the Victorian Government, Australia to I.R.C. and N.E.H.

